# Sensory cheating: adversarial body patterns can fool a convolutional visual system during signaling

**DOI:** 10.1101/326652

**Authors:** Andres Laan, Gonzalo.G de Polavieja

## Abstract

Animals often assess each other by paying special attention to signals, which help to communicate the quality of each individual. When there is a conflict of interest between the signaler and the receiver, then the signaler has an incentive to cheat by producing signals which exaggerate its apparent quality. One opportunity for cheating might be to rely on sensory illusions, but it has been difficult to study sensory cheating because we have lacked quantitative models of complex visual perception. Here we address this problem by taking advantage of recent advances in modeling visual brain areas as convolutional neural networks. Given these models, we use the technique of adversarial perturbations to show how sensory cheating can shape animal appearance while nevertheless resulting in an evolutionarily stable signaling system. In our simulations, animals typically evolve exaggerated color patterns which might be analogous to the evolution of colorful body patterns in guppies.

## Introduction

Convolutional neural networks (CNNs) have recently revolutionized the scientific understanding of image processing and perception [1]. CNNs now form the core component of most modern image recognition software and are routinely used as data analysis tools across many domains. Unlike many previous generations of machine learning models, CNNs are unique because they consist of neuron-like elements and they may thus be viewed as candidate models for explaining the workings of biological visual systems as well. Quantitative comparisons between neural recordings and CNNs have indeed found a close resemblance between neural activity patterns inside CNNs and the mammalian visual cortex [2, 3].

Improved quantitative models of visual perception may provide new ways to theoretically analyze previously intractable problems. Here we use CNN models to study the evolutionary stability of signaling in the presence of conflicts of interest [4]. Our focus will be on the paradigmatic example of aggressive signaling. During aggression, fighters display signals intended to induce their opponent to surrender without a fight [5]. Typically, the individual who is of inferior fighting quality will be scared away by the higher quality individual because the higher quality individual can afford to produce more intense signals. Note that weak individuals theoretically have the option to cheat by somehow producing a more intense signal than the stronger opponent. They are also motivated to do so because successful cheating would lead to easy access to mates and resources. The puzzle of signaling is to explain why cheating does not occur despite strong incentives to do so [4].

Classical models emphasize that such cheating cannot evolve if more intense signals carry with them a greater cost of production, which only high quality individuals are able to bare [6]. This is the standard argument invoked to explain phenomena like the large and uneconomical eyes of the stalk-eyed flies or the peacock’s conspicuous tails[7].

However, the standard explanation can only be part of the answer, because it does not examine stability against sensory cheating. Sensory cheating entails a reshaping of the signal into a form which would make it appear more intense than it really is to the senses of the receiver. The animal would essentially use its own body as a canvas on which to craft a visual illusion. For example, a courting animal might modify its body pigmentation pattern to enhance its apparent height through the use of oriented vertical stripes [8] and subsequently reap the benefits of the illusion through enhanced mating success.

Many animals harness visual illusions in various contexts like camouflage, escape and predator deterrence [9]. A paradigmatic case concerns animals that display large false eyes in order to appear more threatening to predators [10, 11]. Similar phenomena have been reported in the context of signaling as well. Bower birds, for example, are known to actively shape the visual environments of their mates to improve their own mating success [12, 13]. It is therefore not inconceivable that in some species evolution might also shape body patterns so as to trick the sensory systems of the receivers.

In order to quantitatively study the process of sensory cheating, we study a signaling contest where the variable being estimated is body size. We use body size as our variable because larger animals typically win aggressive signaling contests and many animals actively display during conflict to signal their size [5]. We train CNNs to estimate the sizes of model birds placed in natural images. We then let the body pattern of the birds evolve in order to fool the networks and we analyze the emergent dynamics to see if the signaling system remains reliable throughout the process [14].

## Results

Our study considers aggressive contests, where two individuals meet, assess each other’s size and the smaller individual subsequently retreats. Under this scenario, any individual can improve its fitness if it can somehow modify its appearance to appear larger than it really is to the perceptual system of other animals.

To analyze this scenario, we first require a model of the size estimation perceptual system. We therefore compiled a catalog of natural images wherein we placed birds of various sizes. Then we trained a CNN to estimate the size of the bird in each image. After that, we let the birds evolve their appearance in ways that fooled the networks’ perception.

We began by compiling a catalog of 4000 100 by 100 colored natural images. The raw images were downloaded from the natural scene statistics database [15] and 100 by 100 patches were extracted from the first 10 images in the database. We created ten copies of each image and then we placed inside these images the image of a model bird (**Figure 1** left panel). In order to model natural variability in bird appearance, the bird varied in height between 20 and 40 pixels, in rotation between -90 and 90 degrees and its location in the image was also sampled randomly. Further, a different sample of random noise was added to the body of the bird for each image and its intensity was also varied. We thus created a highly variable and non-trivially structured set of 40 000 images whose complexity was designed to mimic the complexity of the natural environment. Sample images of the resulting catalog can be seen in **Figure 1** left panel.

Next we trained a four-layered CNN to predict the size of the bird in each image (see Supplementary methods). Training the CNNs by gradient descent resulted in good predictive accuracy on both the training and the test set (**Figure 1** right panels). The CNNs were thus able to solve the task of separating the birds from the backgrounds and measuring the size of the bird while ignoring irrelevant features like variation in orientation and intensity.

**Figure. 1.**
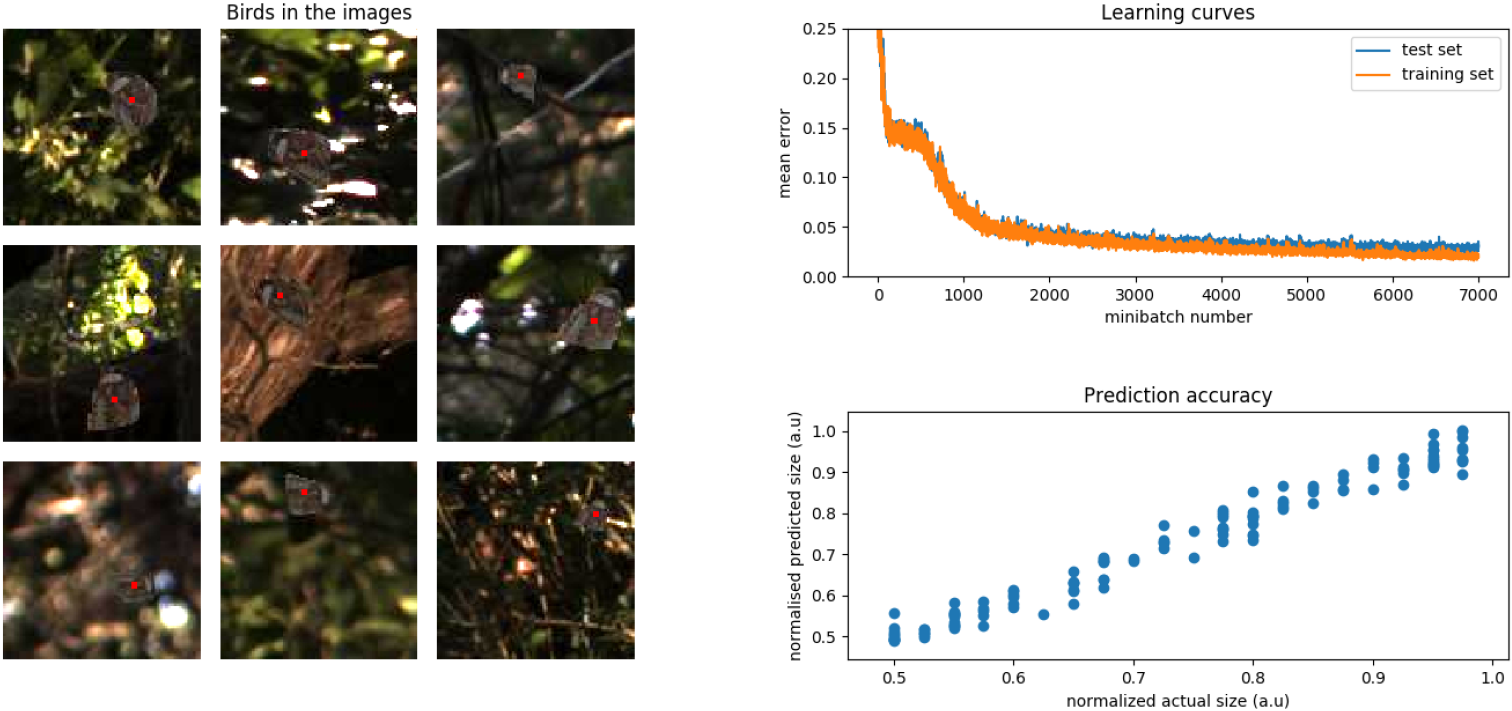
Training the network. Left panel: sample images from the catalog (the center of each bird is marked with a red dot for ease of viewing, the dots were not present in the images on which the CNN was trained). Top right panel: learning curves for the training and test set (batch size was 128 images). Bottom right: correspondence between the ground truth and the output of the trained network.

In order to model the evolution of body patterns, we adapted the technique used to find adversarial examples in artificial neural networks [16]. Briefly, we took three bird images of size 20, 30 and 40 pixels (small, medium, large) and for each image, we calculated the gradient of the output of the network with respect to each pixel of each image. This computation involved estimating the average gradient by taking a sample mean across many backgrounds, orientations and illumination levels (see **Supplementary methods**). Then we performed gradient ascent to make the birds appear progressively larger over each iteration. The evolution of the largest bird’s appearance can be seen in **Figure 2** on the left and the evolution of apparent size is depicted in **Figure 2** right panel.

The birds increase in size by accentuating their edges and decreasing the intensity of the center. They also evolve towards displaying unusual color patterns which are not encountered in the training set. It is noted that though the small, medium and large birds all considerably increase in apparent size, the relative ranking of the sizes of the three birds remains stable throughout evolution and signaling thus remains reliable [14]. Reliability may be conserved because larger individuals are able to cheat more than smaller individuals, because they have more body pixels which they can manipulate. This may help larger individuals maintain their advantage over time.

**Figure. 2.**
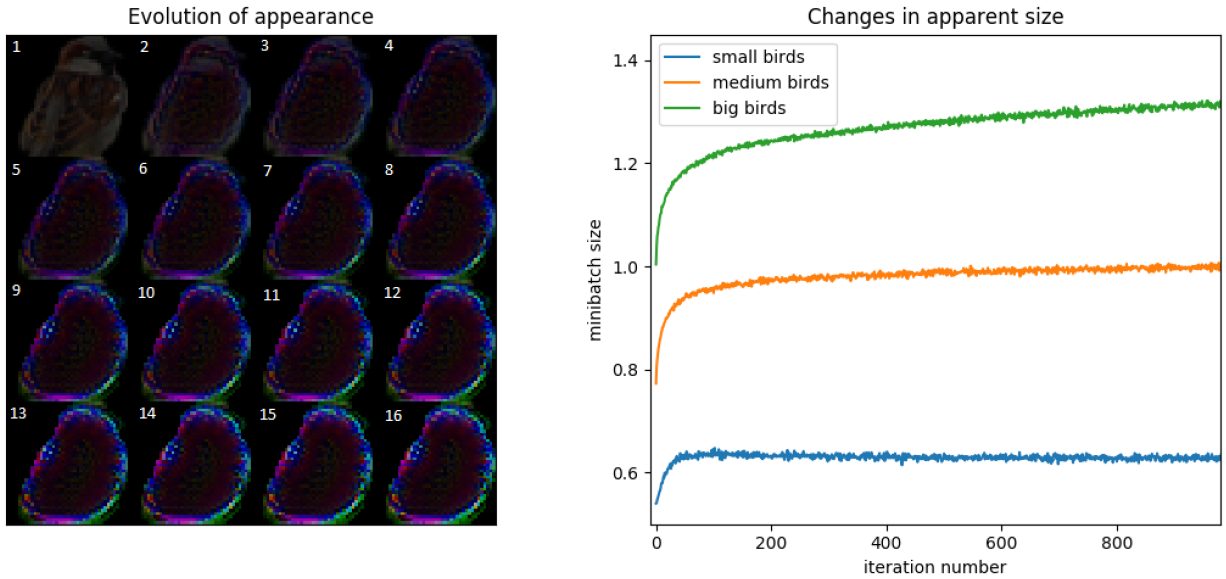
Evolution of apparent size. Left panel: changes in the appearance of the largest bird over time. Time is marked by numbers on the panels, each panel is separated from its predecessor by 60 iterations. Right panel: evolution of apparent size for the small, medium and large bird. Note that all birds considerably increase in size as time progresses but the relative ranking of the sizes nevertheless remains stable.

To establish the suitability of our methods for the study of biological signaling, we further tested whether our conclusions were robust to variation. In biological systems, the cheaters may need to be able to fool multiple networks, since individual brains are known to vary [17]. Though most brains are expected to produce similar outputs for similar inputs, they may be achieving this feat in slightly different ways because internal connections will vary somewhat due to factors like variability in brain development and early visual environment. One way to simulate the variability would be to train many different neural networks from different initial weight values and with a different sequence of training examples.

We implemented this differential training process for five neural networks. We then evolved a bird against the first network and then examined how the findings generalized when the resulting body patterns were shown to the the four other networks. We found that examples developed against one network typically generalized to the other four networks (**Figure 3** top left). We further found that this conclusion held even if we changed the internal architecture of the network when we showed birds evolved against a network using a relu non-linearity to a network using a hyperbolic tangent non-linearity as shown in **Figure S1** left (hyperbolic tangent non-linearities could be viewed as more biologically realistic due to its saturating behavior which more closely mimics neuronal biophysics [18]). As expected, this conclusion did not hold true when the images were shown to an untrained neural network (**Figure S1** right).

**Figure. 3.**
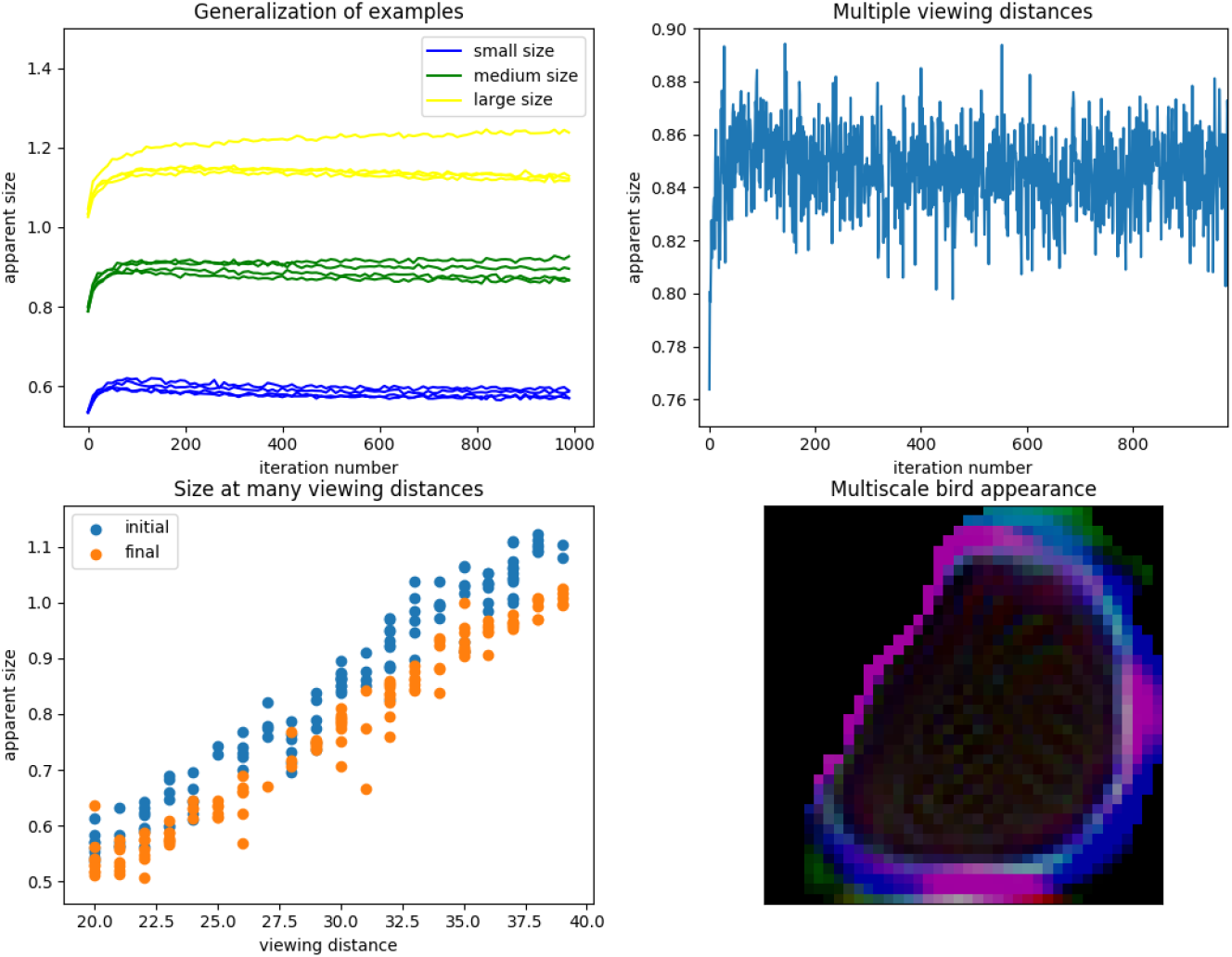
Evolution of apparent size. Top left: Birds evolved against one network are able to fool other networks which they did not encounter during evolution (each trace represents a separate network). Top right: evolution of mean apparent size when viewing distance varies. Bottom left: perceived size versus viewing distance at the beginning (orange) and end (blue) of evolution. The blue dots tend to lie above the red dots for all viewing distances indicating the ability of the mutant to robustly fool the CNN under many conditions. Bottom right: the final appearance of the large bird that fools CNNs at all distance.

We also tested whether our results are robust for all viewing distances. When viewed from an identical distance, smaller animals should always occupy a smaller area on the retina than larger animals. However, signaling displays are often complex spatial maneuvers during which the viewing dis-tance may vary [19]. A good quality visual system would presumably be able to distinguish between a big animal that is far away from a small animal that is close even if both images occupy similar sizes on its retina. Based on these concerns, we trained adversarial examples to be robust against variations in viewing angle (see **Supplementary methods**). Our system was able to find bird pigmentation perturbations which appeared larger than the original bird image at all viewing distances (**Figure 3** top right and bottom panels). We conclude that sensory cheating should be possible even against a visual system which integrates information about image size with information about inter-animal distances.

## Discussion

We have demonstrated how convolutional neural networks could be applied to the study of the evolutionary stability of signaling. We suggest that future studies which examine the stability of signaling models should augment traditional low-dimensional game theory analysis with a high-dimensional analysis of signal form and natural image statistics [4, 6]. Our work shows that unless sensory cheating is ruled out, the stability of any equilibrium cannot be guaranteed.

Our approach made use of the technique of adversarial perturbations, which was originally developed as a method to find small perturbations that will cause machine classifiers to mis-classify an image [16]. Although these perturbations were initially believed to be relevant only in the context of artificial intelligence, recent research indicates that adversarial examples have a limited ability to confuse human observers as well [20]. Our study indicates that adversarial examples may also have further biological relevance in the evolution of signaling and body patterns. Future work could also attempt to apply these techniques to the study of segmentation systems and the evolution of camouflage [21].

Our work is not the first to recognize the usefulness of explicit cognitive models for the study of evolution. Pioneering theoretical work by Enquist and others used artificial neural networks like the multi-layer perceptron as a tool in the theoretical study of evolution [22]. This early work was of limited applicability because slow computers did not allow these systems to be trained on complex real world tasks. With the availability of fast modern hardware, it should become increasingly easy to design and probe the function of complex pattern recognition systems through an evolutionary lens.

One of the empirical findings of our work was that in the later stages of evolution, the model birds evolved to display unusual colors. This is an outcome that likely occurs because adaptive systems are typically tuned to work accurately only on their training domain as they do not face selective pressure to correctly analyze out of domain signals. Since bright and pure colors lie outside the typical statistics of natural images, it is not surprising that these signals turned out to be effective at driving spurious signaling activity in the networks. These findings may have some parallels with the evolution of bright body colors in Trinidadian guppies [23]. When relived from predation pressure, Trinidadian guppies evolve to display bright colors for the purposes of increasing their attractiveness to potential mates. It may be the case that these bright body patterns function partly as adversarial examples or hyper-stimuli that are particularly effective at driving the activity of the sexual quality assessment network of the fish brain.

Our work focused on the evolution of body patterns without considering the simultaneous evolution of the neural network used for assessment. We made this modeling choice because signaling equilibriums may be understood as Bourgeois strategies (where the asymmetry happens to be correlated but need not remain so throughout evolution) and no individual has an incentive to deviate from consensus assessments [24]. Since our modeling finds that signaling remains reliable, it could also serve as a useful model for scenarios where body pattern evolution is for some reason much more rapid than the evolution of the assessment network. For more complex scenarios like the study of sexual selection, this approximation may not remain valid and future work must find ways to extend our methods to take into account the aforementioned complexities [14].

Finally, it will be interesting to study if certain body patterns or brain architectures are less vulnerable to cheating. It might be expected that pure bright colors which are already unusual for a given environment and easy to separate from the background might be rather immune to cheating. Also, there may be other neural networks which utilize movement information or do a more complex segmentation that will prove more difficult to hack. Future work will need to explore these issues in more extensive detail.

## Supplementary materials

### Supplementary figures

**Figure. 4.**
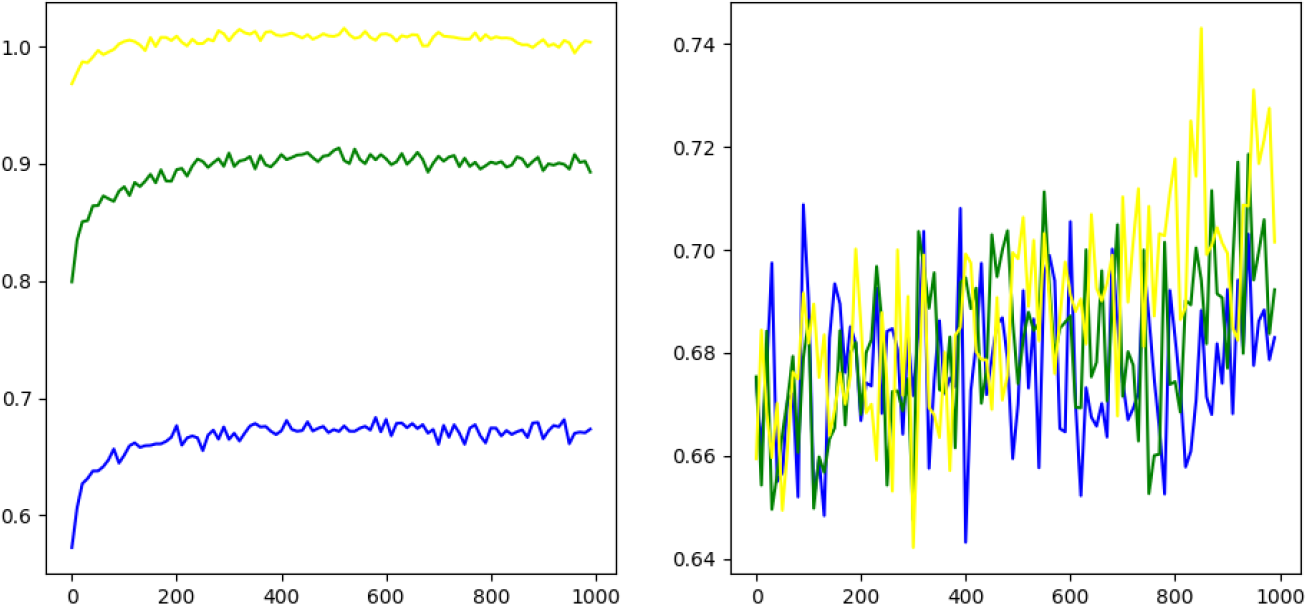
Generalization of examples. Left: Birds evolved against a relu network shown to a hyperbolic tangent network. Right: Birds evolved against a trained relu network shown to an incompletely trained (100 instead of 7000 mini-batches) relu network.

## Supplementary methods

### Details on neural networks

We used 4 convolutional layers with relu non-linearities, each followed by a 2-by-2 max pooling layer. All layers used 5x5 filters. Filter numbers by layer were 32, 64, 64, 64. The fully connected layer used 512 neurons. We trained the network using gradient descent on the mean squared error loss function with the Adam optimizer using a learning rate of 10^−4^ with mini-batches of size 128. The training set consisted of 35 000 images from which mini-batches were sampled randomly. For the relu networks training process used 7000 mini-batches. For the tanh non-linearity training took 200 000 mini-batches. Training was implemented in Tensorflow.

### Adversarial examples

During evolution, birds will evolve towards greater apparent size. The image of a bird can be regarded as a set of pixels. To predict how the birds will evolve, we must predict which pixel changes would increase the expected apparent size of the bird. In other words, we must predict the gradient of the expected apparent size with respect to each pixel of the bird. In order to estimate the expected value of the gradient, we must average over all possible locations, orientations, backgrounds, noise perturbations, etc. We calculate the estimate using Monte Carlo sampling. We first embed the bird in 128 images, whose orientation, location, background, etc statistics are sampled from the same distribution as was used for generating the training set. For each image, we use standard Tensorflow procedures to estimate the gradient of the output (the estimated size) with respect to all the image pixels. Then we back-transform this gradient onto the bird image template by shifting and rotating the image such that the birds in all the images will line up exactly. The gradient estimate is the sample mean of these back-transformed gradients. Then we add the learning-rate weighted gradient onto the bird images to obtain a new bird image and repeat the procedure again. The same procedure was used to in the viewing distance invariant scenario, but there we also added an extra image scaling step to the back-transformation step to compensate for the fact that the size of the bird in each images varied depending on the viewing distance (the viewing distance was sampled uniformly at random between 20 and 40 units). Further details on the code are available from the authors on request and all code will be deposited at a public repository after publication of the manuscript.

## References

[1] Krizhevsky A, Sutskever I, Hinton GE. Imagenet classification with deep convolutional neural networks. In: Advances in neural information processing systems; 2012. p. 1097–1105.

[2] Khaligh-Razavi SM, Kriegeskorte N. Deep supervised, but not unsupervised, models may explain IT cortical representation. PLoS computational biology. 2014;10(11):e1003915.

[3] Kriegeskorte N. Deep neural networks: a new framework for modeling biological vision and brain information processing. Annual review of vision science. 2015;1:417–446.

[4] Smith JM, Harper D. Animal signals. Oxford University Press; 2003.

[5] Arnott G, Elwood RW. Assessment of fighting ability in animal contests. Animal Behaviour. 2009;77(5):991–1004.

[6] Grafen A. Biological signals as handicaps. Journal of theoretical biology. 1990;144(4):517–546.

[7] Emlen DJ.Animal weapons: the evolution of battle. Henry Holt and Company; 2014.

[8] Thompson P, Mikellidou K.Applying the Helmholtz illusion to fashion: Horizontal stripes won’t make you look fatter. i-Perception. 2011;2(1):99–76.

[9] Kelley LA, Kelley JL. Animal visual illusion and confusion: the importance of a perceptual perspective. Behavioral Ecology. 2013;25(3):450–463.

[10] De Bona S, Valkonen JK, Lopez-Sepulcre A, Mappes J. Predator mimicry, not conspicuousness, explains the efficacy of butterfly eyespots. Proc R Soc B. 2015;282(1806):20150202.

[11] Stevens M, Hardman CJ, Stubbins CL. Conspicuousness, not eye mimicry, makes eyespots effective antipredator signals. Behavioral Ecology. 2008;19(3):525–531.

[12] Endler JA, Endler LC, Doerr NR. Great bowerbirds create theaters with forced perspective when seen by their audience. Current Biology. 2010;20(18):1679–1684.

[13] Kelley LA, Endler JA. Illusions promote mating success in great bowerbirds. Science. 2012;335(6066):335–338.

[14] Searcy WA, Nowicki S. The evolution of animal communication: reliability and deception in signaling systems. Princeton University Press; 2005.

[15] Geisler WS, Perry JS. Statistics for optimal point prediction in natural images. Journal of Vision. 2011;11(12):14–14.

[16] Szegedy C, Zaremba W, Sutskever I, Bruna J, Erhan D, Goodfellow I, et al. Intriguing properties of neural networks. arXiv preprint arXiv:13126199. 2013;.

[17] Maguire EA, Gadian DG, Johnsrude IS, Good CD, Ashburner J, Frack-owiak RS, et al. Navigation-related structural change in the hippocampi of taxi drivers. Proceedings of the National Academy of Sciences. 2000;97(8):4398–4403.

[18] Byrne JH, Heidelberger R, Waxham MN. From molecules to networks: an introduction to cellular and molecular neuroscience. Academic Press; 2014.

[19] Laan A, Iglesias M, de Polavieja G. Opponent assessment and conflict resolution through mutual motor coordination. bioRxiv. 2017;p. 208918.

[20] Elsayed GF, Shankar S, Cheung B, Papernot N, Kurakin A, Goodfellow I, et al. Adversarial Examples that Fool both Human and Computer Vision. arXiv preprint arXiv:180208195. 2018;.

[21] Stevens M, Merilaita S. Animal camouflage: current issues and new perspectives. Philosophical Transactions of the Royal Society B: Biological Sciences. 2009;364(1516):423–427.

[22] Enquist M, Ghirlanda S. Neural networks and animal behavior. Princeton University Press; 2013.

[23] Endler JA. Natural selection on color patterns in Poecilia reticulata. Evolution. 1980;34(1):76–91.

[24] Sherratt TN, Mesterton-Gibbons M. The evolution of respect for property. Journal of evolutionary biology. 2015;28(6):1185–1202.

